# CaMutQC: An R Package for Integrative Quality Control of Cancer Somatic Mutations

**DOI:** 10.1101/2024.08.12.606123

**Authors:** Xin Wang, Tengjia Jiang, Ao Shen, Yaru Chen, Yanqing Zhou, Jie Liu, Shuhan Zhao, Shifu Chen, Jian Ren, Qi Zhao

**Affiliations:** State Key Laboratory of Oncology in South China, Guangdong Key Laboratory of Nasopharyngeal Carcinoma Diagnosis and Therapy, Guangdong Provincial Clinical Research Center for Cancer, Sun Yat-sen University Cancer Center, Guangzhou 510060, China; Department of Bioinformatics, HaploX Biotechnology, Shenzhen, 518057, China

## Abstract

The quality control and filtration of cancer somatic mutations (CAMs), including the elimination of false positives resulting from technical bias and the selection of key mutation candidates, is crucial for downstream analysis in cancer genomics. Due to diverse needs and the absence of standardized filtering criteria, the filtering strategies employed vary from study to study, often leading to reduced efficiency, accuracy, and comparability across similar analyses. Here we present CaMutQC, a heuristic quality control and soft-filtering R/Bioconductor package for CAMs. With CaMutQC, the removal of false positives, selection of potential mutation candidates, and Tumor Mutation Burden estimation can be executed in a single line of code, using default or customized parameters. A filter report and a code log generated after the filtration process assist with recording and comparison. The application of CaMutQC on a Whole-exome Sequencing (WES) benchmark dataset demonstrated its impressive capability by eliminating 85.55% of false positive mutations while retaining 90.72% of true positive mutations. Additionally, an extra 11.56% of true positive mutations were recused by the union strategy embedded in CaMutQC. CaMutQC is now available through Bioconductor at https://bioconductor.org/packages/CaMutQC/ under the GPL v3 license, and it will be updated regularly to incorporate top filtration strategy and parameter sets shared within the community.

## Introduction

Driven by the continuous accumulation of somatic mutations, tumorigenesis is generally regarded as an evolutionary process involving genetic mutation and natural selection. Approximately 90% of cancer genes exhibit somatic mutations^1^, with diverse patterns manifesting across different cancer lineages^2^ and among individuals with the same cancer type. This variability significantly contributes to genetic heterogeneity in cancer, underscoring the importance of analyzing cancer somatic mutations (CAMs) for developing targeted and personalized treatments.

With advances in sequencing technologies, particularly the Next-generation Sequencing (NGS), significant progress has been made in detecting somatic mutation and identifying mutational patterns across various cancer types^3^. Annotation databases have further broadened our understanding of CAMs on a larger scale. However, artifacts might be introduced due to technique issues during some processes: DNA fixation^4,5^, the fragmentation and chemical modifications occurring during the Formalin-Fixed Paraffin-Embedded (FFPE) process^6^, as well as DNA oxidation and the formation of 8-oxoguanine (8-oxoG) lesions^7^. How to remove these artifacts accurately and efficiently is of great significance when processing CAMs. Although tools like MuTect2^8^ and VarScan2^9^ have been developed to detect cancer somatic mutations, limitations exist when applying them to real-world data: (1) parameters and callers used in different studies usually vary, making it difficult to conduct cross-study research (2) FPs and false negatives (FNs) can be significant when using single caller, and rescue cannot be performed through traditional calling methods. While R-based tools like vcfR^10^ and isma^11^ can read, visualize, and compare somatic mutations, none support comprehensive and integrative cancer-specific somatic mutation filtration.

To address the need for systemic and customized CAM filtration, we developed CaMutQC, a heuristic cancer somatic mutation quality control and filtration software. CaMutQC labels every mutation with distinct tags corresponding to various filtering strategies. It also generates a vivid and detailed filter report after a complete filtration and screening, offering transparent insights into filtration outcomes. We have built a community where users are encouraged to share their strategies and parameters, promoting collaboration and communication in CAM quality control. Our commitment to continuous improvement includes incorporating top-rated strategies shared in the community into CaMutQC, ensuring easy accessibility to an evolving set of effective strategies. Furthermore, we propose an updated union strategy for the quality control of CAMs: taking the union of CaMutQC-filtered cancer mutations from multiple variant callers. We believe this union strategy has great potential to rescue true positive mutations (TP) while eliminating more FPs simultaneously, providing a superior alternative to relying on a single variant caller.

## Materials and Methods

### Data collection and preprocessing

We applied CaMutQC on two WES datasets, WES_EA_1 and WES_FD_2, along with one WGS dataset WGS_EA_1, all the datasets were collected from an NCBI study^12^. Starting from paired tumor-normal BAM files, three variant callers --- MuTect2, VarScan2 and MuSE^13^ --- were run on all three datasets with default parameters, and the internal filters of variant callers were applied if applicable. For two WES datasets, we defined target regions by intersecting high-confidence regions with WES sequencing regions, while setting the high-confidence regions as target regions for the WGS dataset. Mutational calls were annotated before and after internal filtrations using Variant Effect Predictor^14^ (VEP) (v110).

### Convert VCF to Mutation Annotation Format (MAF)

Currently, vcf2maf^15^ is widely used to convert VCF to MAF, but it requires a Linux environment with Perl installed and does not accept VEP-annotated VCF files. The *vcfToMAF* function in CaMutQC enables easy conversion with a single line of code while retaining as much information as possible for subsequent steps.

### Common filtering strategies

CaMutQC incorporates numerous widely adopted filtration strategies, encompassing potential artifact filtration and candidate mutation selection, with default parameters collected from previous studies (Table 1). Notably, CaMutQC does not directly exclude CAMs that do not meet filtration criteria but instead labels them with distinct flags, providing additional information for further analyses. As a special feature of CaMutQC, this soft-filtering setting ensures that no valuable information is discarded, empowering users with enhanced insights and flexibility in refining their analyses.

**Table 1.**
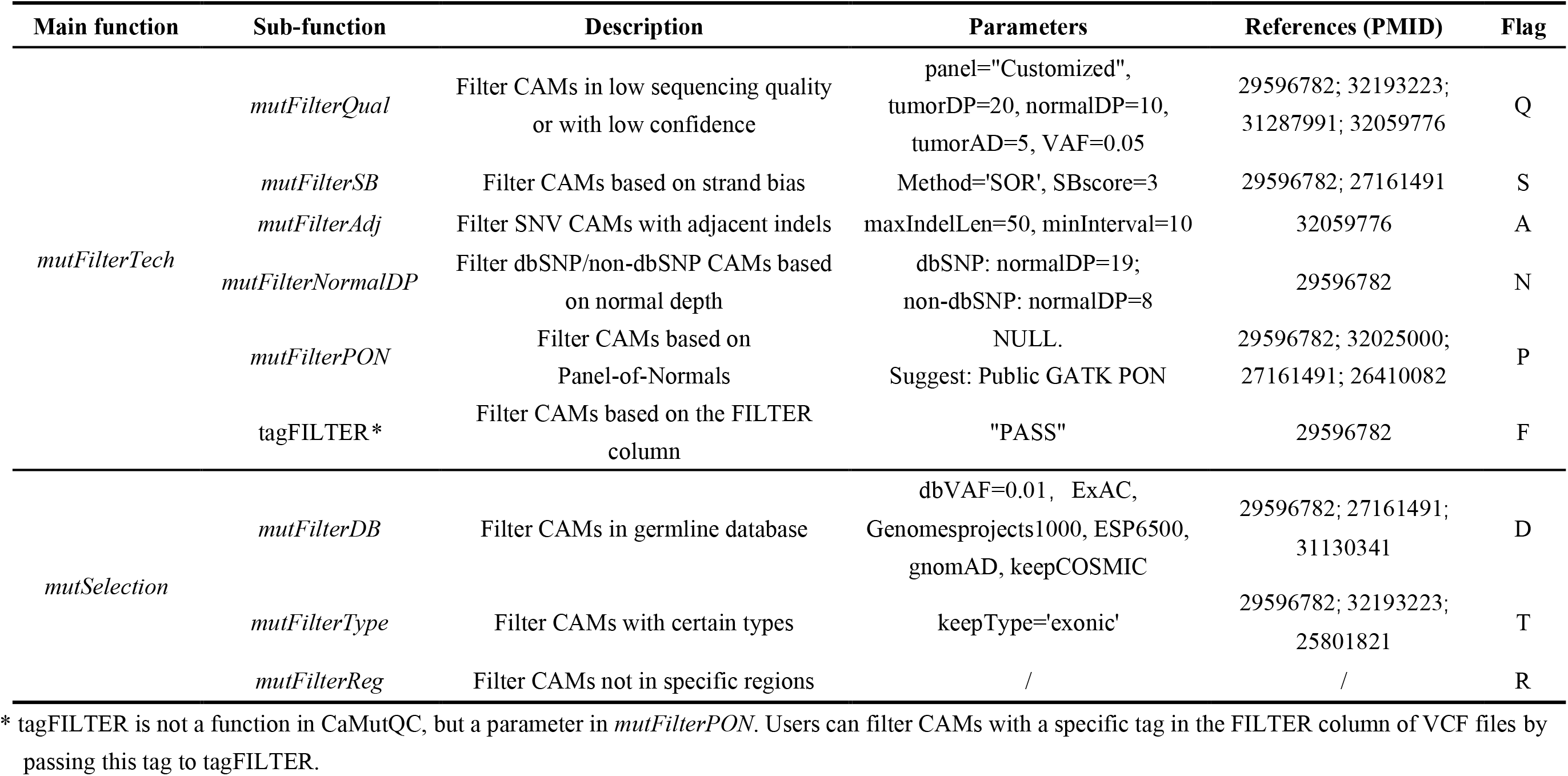
Main functions, sub-functions in CaMutQC, and their description, parameters, and tags.

| Main function | Sub-function | Description | Parameters | References (PMID) | Flag |
| --- | --- | --- | --- | --- | --- |
| *mutFilterTech* | *mutFilterQual* | Filter CAMs in low sequencing quality or with low confidence | panel="Customized",<br>tumorDP=20, normalDP=10,<br>tumorAD=5, VAF=0.05 | 29596782; 32193223;<br>31287991; 32059776 | Q |
|  | *mutFilterSB* | Filter CAMs based on strand bias | Method='SOR', SBscore=3 | 29596782; 27161491 | S |
|  | *mutFilterAdj* | Filter SNV CAMs with adjacent indels | maxIndelLen=50, minInterval=10 | 32059776 | A |
|  | *mutFilterNormalDP* | Filter dbSNP/non-dbSNP CAMs based on normal depth | dbSNP: normalDP=19;<br>non-dbSNP: normalDP=8 | 29596782 | N |
|  | *mutFilterPON* | Filter CAMs based on Panel-of-Normals | NULL.<br>Suggest: Public GATK PON | 29596782; 32025000;<br>27161491; 26410082 | P |
|  | tagFILTER* | Filter CAMs based on the FILTER column | "PASS" | 29596782 | F |
| *mutSelection* | *mutFilterDB* | Filter CAMs in germline database | dbVAF=0.01, ExAC,<br>Genomesprojects1000, ESP6500,<br>gnomAD, keepCOSMIC | 29596782; 27161491;<br>31130341 | D |
|  | *mutFilterType* | Filter CAMs with certain types | keepType='exonic' | 29596782; 32193223;<br>25801821 | T |
|  | *mutFilterReg* | Filter CAMs not in specific regions | / | / | R |

\* tagFILTER is not a function in CaMutQC, but a parameter in *mutFilterPON*. Users can filter CAMs with a specific tag in the FILTER column of VCF files by passing this tag to tagFILTER.

To **remove false positives and germline mutations**, CaMutQC uses critical sequencing quality indicators such as tumor depth (DP), allele depth (AD), and Variant Allele Frequency (VAF). The *mutFilterSB* function within CaMutQC allows users to apply Strand Bias filtration with a default cutoff of 3, measured through the StrandOddsRatio^8^ approach or Fisher’s Exact Test^16^. For germline mutations, CaMutQC can filter out mutations included in the Panel-of-Normal (PON) dataset provided by users and apply different thresholds on normal depth for dbSNP^17^/non-dbSNP mutations. Additionally, SNVs occurring within 10 bps of any INDEL are also labeled as potential FPs by default. Recognizing the superior performance achieved by combining calls from multiple callers^18,19^, CaMutQC introduces an updated union strategy: passing calls from multiple callers first into CaMutQC and taking union afterward. We believe this approach will further elevate performance by rescuing more true positive mutations while effectively eliminating false positives.

To **identify CAMs likely associated with tumorigenesis** from a filtered pool, CaMutQC employs specific criteria. By default, CAMs that either do not exist in germline databases like ExAC^20^ and ESP6500^21^ or exist but exhibit a Variant Allele Frequency (VAF) lower than 0.01, are retained. This stringent filtration process ensures the exclusion of common germline variants while prioritizing those with higher VAFs that may indicate somatic mutations associated with tumorigenesis. In addition to filtering mutations outside of user-provided target region(s), CaMutQC also labels non-exonic CAMs by default. This approach ensures the selected CAMs are more likely to be associated with tumorigenesis, as exonic mutations often matter more in tumorigenesis than non-exonic ones.

### Customized filtering strategies

Recognizing the diverse needs in cancer genomics research, filtering strategies applied to CAMs often exhibit variability across studies and cancer types. CaMutQC addresses this need by implementing the *mutFilterCan* and *mutFilterRef* functions, allowing users to apply cancer type-specific and reference-specific strategies, respectively. With these two functions, greater comparability between studies can be achieved by applying the same set of strategies with ease, potentially facilitating the establishment of filtering standards. (Table 2).

**Table 2.**
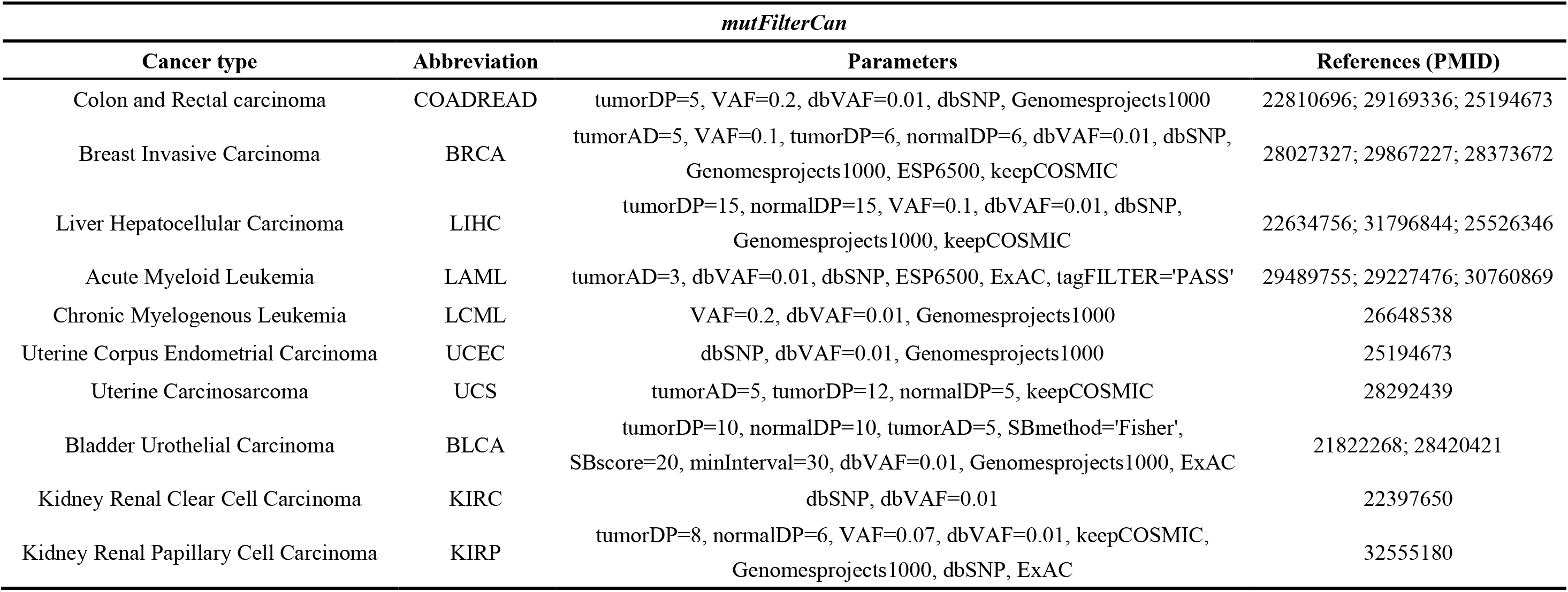
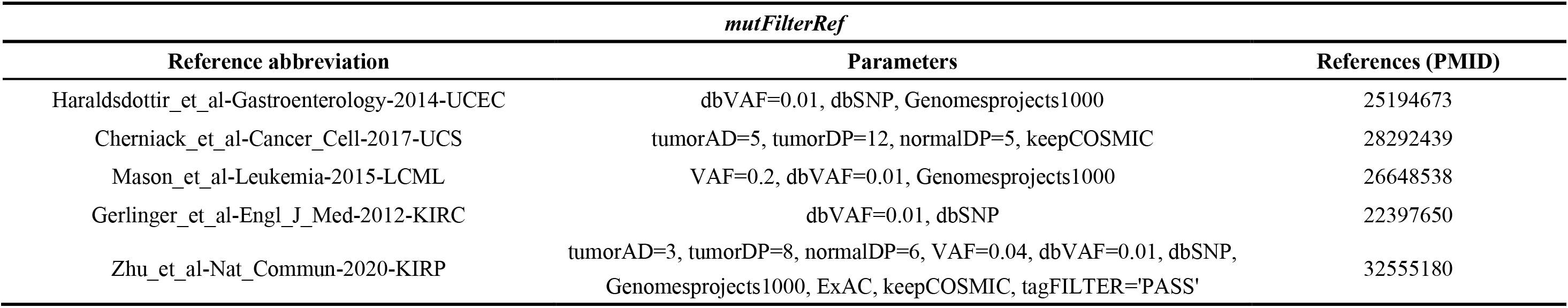
Details of two main functions in CaMutQC, including parameters, references, etc.

### Tumor Mutational Burden (TMB) estimation

TMB quantifies the number of non-synonymous somatic mutations per megabase pair (Mb) within a specific genomic region. In 2015, TMB was first reported to be linked to PD1/PD-L1 cancer immunotherapy^22^ and has been viewed as a critical standard in clinical practice^23,24^. Several assays and panels are available for TMB measurement, and CaMutQC supports four of them: FoundationOne^25^, MSK-IMPACT (3 versions)^26^, Pan-cancer panel^27^, and WES. By default, TMB is calculated to align with the MSK-IMPACT-v3 assay, but users can apply their methods under the “Customized” mode. It is worth noting that CaMutQC utilizes BED region files generated solely from the coding sequences (CDS) of panel genes. Although these regions may not be the exact region, the TMB estimations provided by CaMutQC serve as a valuable reference.

## Results

### Overview of CaMutQC functions and implementation

As an open-source R/Bioconductor package, CaMutQC takes VEP-annotated VCF file(s) as input. CaMutQC supports 3 callers: MuTect2, VarScan2 and MuSE, and can also operate under multi-sample or multi-caller modes. After transforming VCFs into MAFs, CaMutQC provides a series of default or customized filtrations and selections on CAMs, along with TMB estimations with various assays (Figure 1A). A labeled MAF and a detailed filter report will be generated after a whole mutation filtration or selection section. Moreover, CaMutQC establishes direct connections with downstream tools MesKit^28^ and maftools^29^ (Figure 1B). This integration facilitates seamless transitions between software, enhancing user convenience and the overall utility of CaMutQC in cancer genomics research. It is important to note that additional files may be required by the target software during the transition.

**Figure 1.**
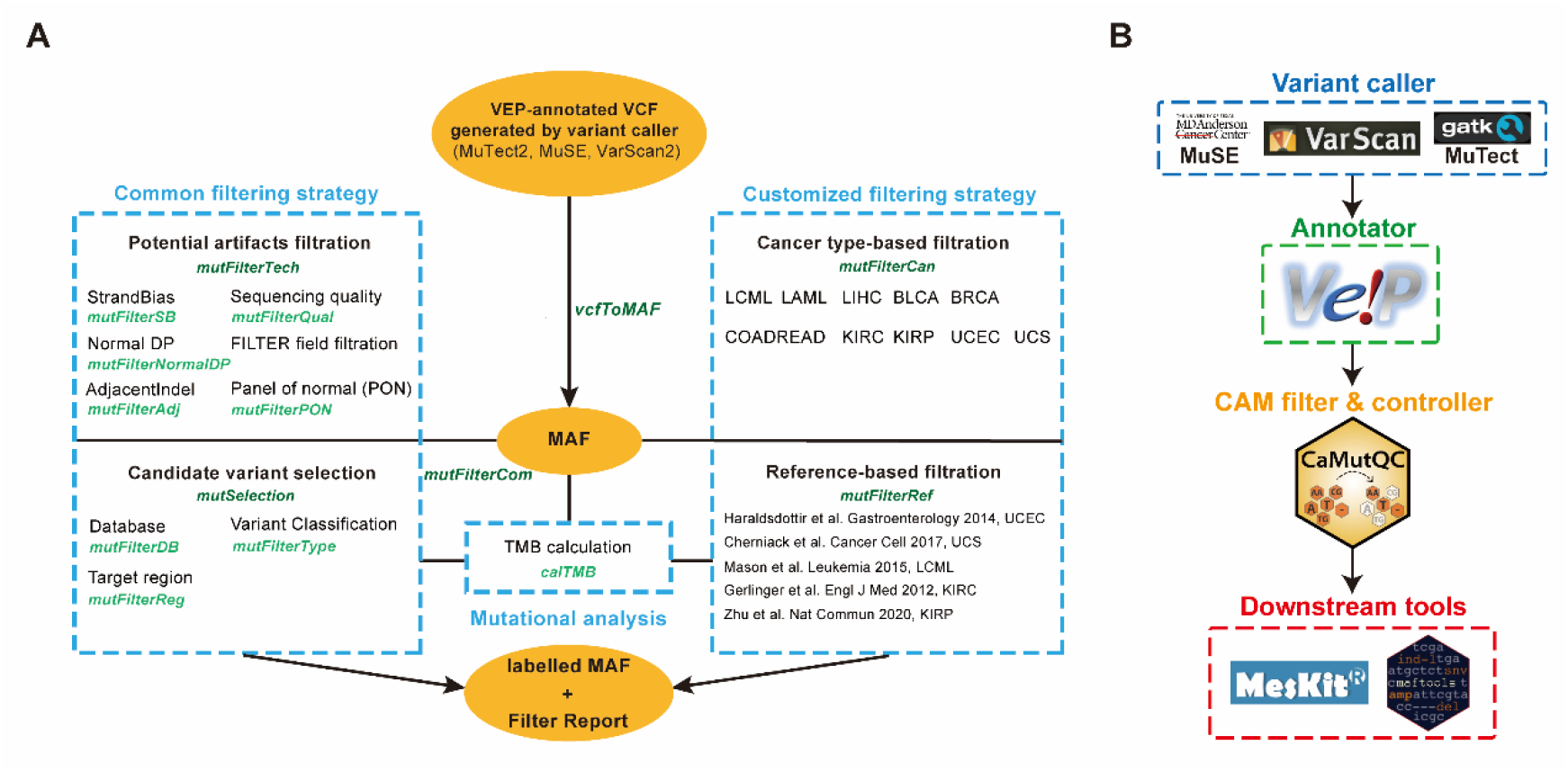
**A**. An overview of CaMutQC. **B**. CaMutQC connects the upstream and downstream of variant calling by taking annotated variant calls, filtering them, and passing them to downstream tools.

### Benchmark results of CaMutQC

Two main functions in CaMutQC, *mutFilterTech*, and *mutSelection*, were benchmarked with default parameters. The PON dataset used during this process was downloaded from the public GATK resources bundle (public Google bucket: gatk-best-practices). To validate *mutFilterTech*, we compared FPs and TPs before and after filtration against the total number of TPs for the two WES datasets. Figure 2 and Table S1 show that CaMutQC efficiently retains most of the TP mutations even when internal filters were not applied, underscoring its stability, consistency, and minimal dependency on other filters. Nevertheless, we focused on internal-filtered mutations in subsequent analyses, given the conventional view of internal filtration as an integral part of a caller. In the WES_EA_1 dataset, with an average sequencing depth of 60x, *mutFilterTech* impressively removed 85.55% of FP single nucleotide polymorphisms (SNPs) while retaining 90.72% of TP SNPs across the three callers on average. The reasons why a higher percentage of FP insertions and deletions (INDELs) failed to be removed might be attributed to two factors: 1) there are significantly fewer TP INDELs in the target region compared to TP SNPs (50 compared to 1159), leading to a much smaller denominator when calculating the percentage; 2) MuSE does not support INDEL calling, and VarScan2 has been found to perform poorly on calling INDELs with shortread data^30^, introducing more uncertainties and potential biases. The impact of sequencing depth on post-calling quality control was evident when comparing results from two WES datasets, where the removal rate for both FP and TP increased in SNPs and INDELs in WES_FD_2 (Figure S1). These findings highlight the crucial role of higher sequencing depth in achieving accurate results. In terms of the WGS dataset, since researchers usually turn on internal filters by default to reduce the number of mutations, we only ran CaMutQC on internal-filtered mutations. Results again show that CaMutQC has great potential in removing FPs, thus improving the accuracy of following analyses (Figure S2).

**Figure 2.**
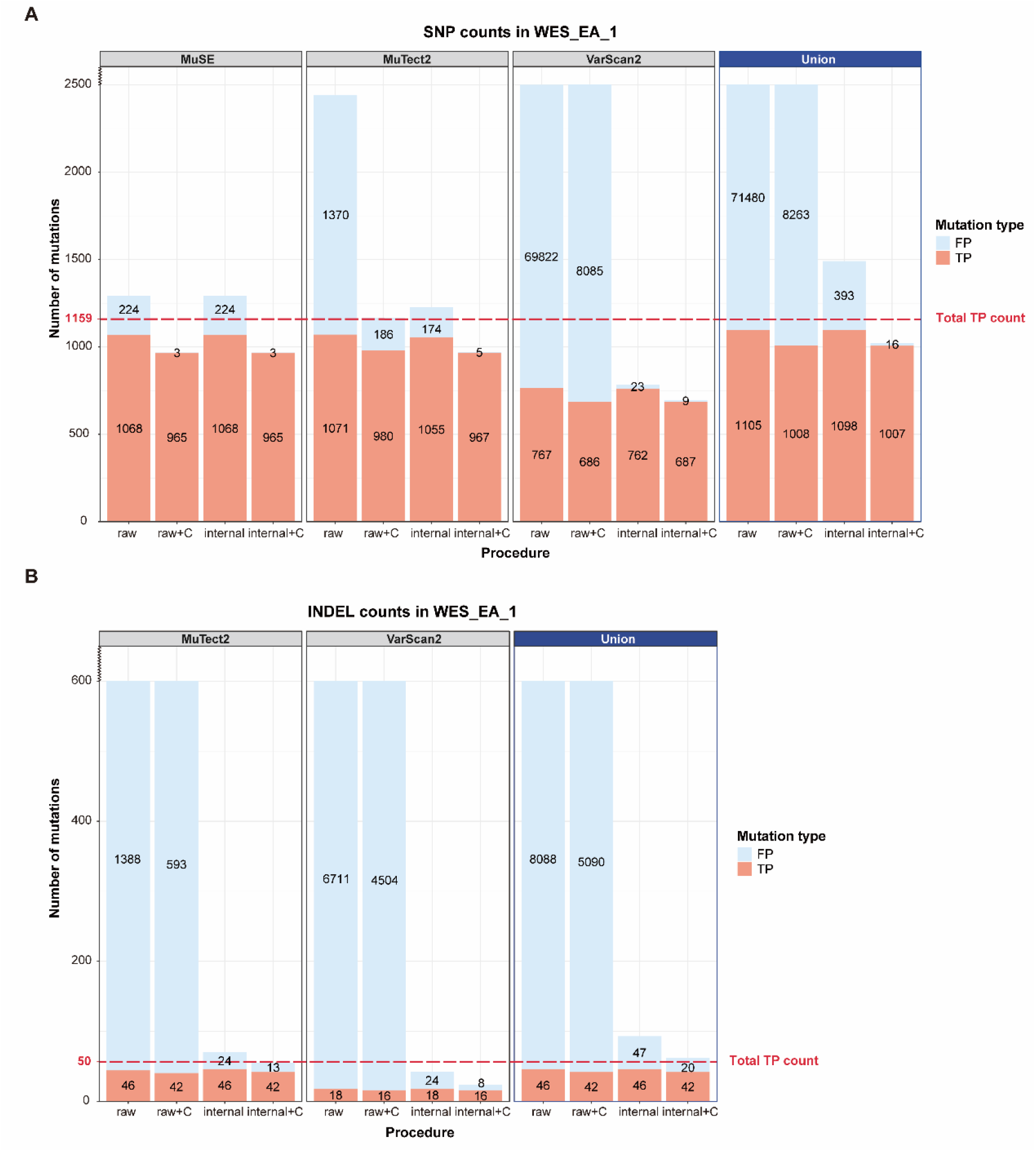
Benchmark results of WES_EA_1. **A**. TP and FP SNP counts after different procedures. The number of FP SNPs has greatly decreased after CaMutQC filtration with/without internal filtration. **B**. TP and FP INDEL counts after different procedures. raw: raw counts with no filtration; C: filtered by CaMutQC (CaMutQC-*mutFilterTech*); internal: processed by callers’ internal filter.

### Rescue TP through updated Union strategy

To validate the proposed union strategy, we took the union of CaMutQC-filtered mutations from MuTect2, MuSE, and VarScan2 (Figure 3). The results on the WES_EA_1 dataset revealed its significant potential in rescuing TP mutations, recalling an additional 18.36% of TP SNP, or 11.56% of total TP SNP counts compared to the single variant caller. Similarly, the union strategy successfully rescued some TP INDELs, further underscoring its effectiveness.

**Figure 3.**
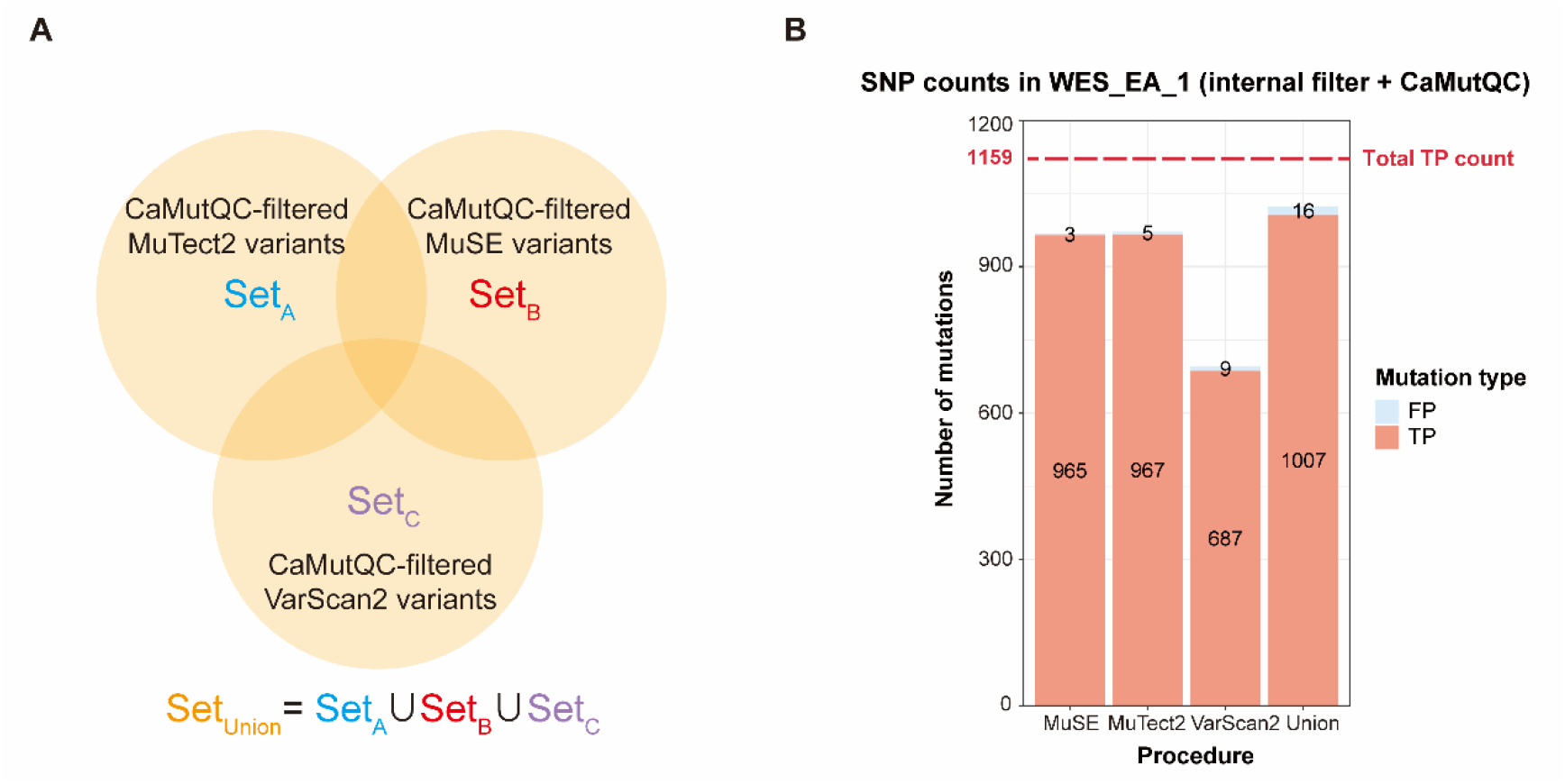
An example of the updated union strategy. **A**. An illustration of the union strategy applied in this study. The union of CaMutQC-filtered (CaMutQC-*mutFilterTech*) variants from three callers, MuTect2, VarScan2, and MuSE, was obtained. **B**. Union results of SNPs in WES_EA_1. The number of TP increases by 18.36% on average among three callers.

### Selection of key somatic mutations

*mutSelection* function, designed to select potential candidate mutations related to tumor development and growth post *mutFilterTech* filtration, also demonstrated its effectiveness. When *mutFilterTech*-filtered mutations from the WGS dataset were passed into *mutSelection*, only 1.35% of SNP and 1.16% of INDEL were selected, where 51.33% of selected SNPs were recorded in COSMIC^31^ (Table S2). This indicates the proficiency of CaMutQC in selecting key mutations while allowing for exploration beyond known candidate mutations.

## Discussion

Somatic mutations play a significant role in oncogenesis, revealing tumor-specific molecular traits crucial for diagnosis and treatment. However, challenges arise in identifying true positive somatic mutations due to biases in sample processing and disparities among sequencing platforms. Recently, the focus has shifted towards calling somatic mutations with high precision and confidence, highlighting the need for a universally acknowledged, accepted, and shared post-calling quality control mechanism within the field.

In clinical practices, researchers expect to have as many somatic mutations as possible to understand tumor progression better and explore potential treatment options. Meanwhile, bioinformatics analysts prioritize the confidence and accuracy of mutations. With CaMutQC, this conflict can be mitigated by reviewing mutation flags and making tailored decisions. Designed specifically for CAMs, CaMutQC supports input variant calling results from three popular callers, performing quality control and filtration based on strategies collected from established studies. Benchmark results highlight CaMutQC’s exceptional performance in flagging false positives, and the soft-filtering concept further enhances its adaptability to diverse needs. A comprehensive filter report and code log improve the user experience and boost community collaboration. In addition, various options for TMB estimation and functions that connect CaMutQC with downstream tools expand its application in research and clinical domains.

As we strive for continuous maintenance and enhancement, CaMutQC will evolve in various aspects. Future updates will include supporting inputs from more variant callers like SomaticSniper^32^ and more annotators like ANNOVAR^33^. Ongoing efforts also target refining and updating filtering strategies. Different thresholds might be established and tested for samples with different sequencing depths since we observed better performance in the WES dataset with higher depth. Moreover, a whitelist with key genes and variants will be introduced in future versions. Inspired by DeepVairant^34^ the integration of machine learning algorithms into the quality control process of CaMutQC holds promise for optimizing performance. Exploring the capabilities of artificial intelligence in this context could provide valuable insights and potentially elevate the efficacy of somatic mutation filtration.

## Availability of Source Code and Requirements

Project name: CaMutQC

Project home page: https://github.com/likelet/CaMutQC

Operating system(s): Platform independent

Programming language: R

Other requirements: R ≥ 4.0

License: GPL-3.0

## Data availability

Public benchmark datasets analyzed in this study, along with reference genome (GRCh38.d1.vd1), high-confidence mutations set (used as TP set in this study), and high-confidence regions, were obtained from NCBI’s FTP site (ftp://ftp-trace.ncbi.nlm.nih.gov/ReferenceSamples/seqc/Somatic_Mutation_WG)^12^.

## Authors’ Contributions

Q.Z. and J.R. conceived the project. T.J. carried out related research and set the filtering parameters. X.W. developed and implemented the methodology. X.W., A.S., J.L., and

S.Z. helped test the software. Y.C., Y.Z., and S.C. helped supervise the project. X.W., and Q.Z. wrote the manuscript. All authors read and approved the final manuscript.

## Acknowledgments

This work is supported by the Young Talents Program of Sun Yat-sen University Cancer Center (YTP-SYSUCC-0033 to Q.Z).

