## Supplemental Figure 1 for "CaMutQC: An R Package for Integrative Quality Control of Cancer Somatic Mutations"

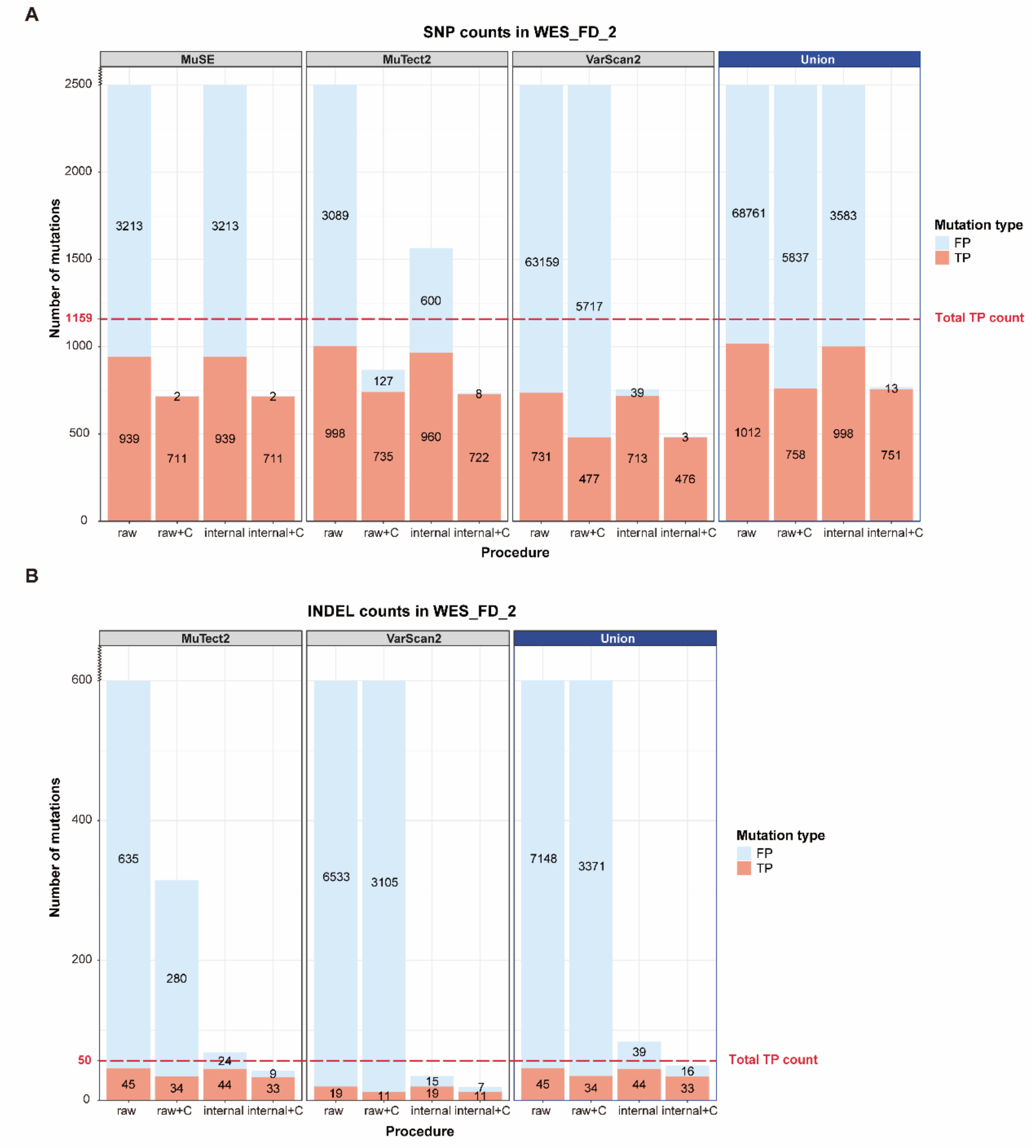


**Supplemental Figure 1.**

Benchmark results of sample WES_FD_2. **A.** TP and FP SNP counts after different procedures. **B.** TP and FP INDEL counts after different procedures. raw: raw counts with no filtration; C: filtered by CaMutQC (CaMutQC-*mutFilterTech*); internal: processed by callers' internal filter.
